# Many ways to make darker flies: Intra- and inter-specific variation in *Drosophila* body pigmentation components

**DOI:** 10.1101/2020.08.26.268615

**Authors:** Elvira Lafuente, Filipa Alves, Jessica G King, Carolina M Peralta, Patrícia Beldade

## Abstract

Body pigmentation is an evolutionarily diversified and ecologically relevant trait that shows variation within and between species, and important roles in animal survival and reproduction. Insect pigmentation, in particular, provides some of the most compelling examples of adaptive evolution and its ecological and genetic bases. Yet, while pigmentation includes multiple aspects of color and color pattern that may vary more or less independently, its study frequently focuses on one single aspect. Here, we develop a method to quantify color and color pattern in *Drosophila* body pigmentation, decomposing thorax and abdominal pigmentation into distinct measurable traits, and we quantify different sources of variation in those traits. For each body part, we measured overall darkness, as well as four other pigmentation properties distinguishing between background color and color of the darker pattern elements that decorate the two body parts. By focusing on two standard *D. melanogaster* laboratory populations, we show that pigmentation components vary and co-vary in different manners depending on sex, genetic background, and developmental temperature. By studying three natural populations of *D. melanogaster* along a latitudinal cline and five other *Drosophila* species, we then show that evolution of lighter or darker bodies can be achieved by changing distinct component traits. Our study underscores the value of detailed phenotyping for a better understanding of phenotypic variation and diversification, and the ecological pressures and genetic mechanisms underlying them.

## INTRODUCTION

Diversity in body coloration provides some of the most compelling examples of adaptive evolution. Insect body coloration, in particular, includes text book cases such as industrial melanism (e.g. van’t Hof et al. 2011; Cook and Saccheri 2013), mimicry (e.g. Mallet & Joron 1999; Nadeau 2016), and clinal variation (e.g. Bastide et al. 2014; Endler et al. 2016). Studies in different species have illustrated the ecological significance of variation in body pigmentation, including visual communication between individuals of the same (e.g. mate attraction and mate choice; e.g. Wiernasz 1995; Guillermo-Ferreira et al. 2014) or different species (e.g. predator avoidance via camouflage or aposematism; e.g. Reichstein et al. 1968; Futahashi and Fujiwara 2008; van Bergen and Beldade 2019), as well as thermoregulation (e.g. Rajpurohit et al. 2008; Sibilia et al. 2018). Moreover, insect pigmentation is tightly associated with various other traits that are closely related to fitness (see Wittkopp and Beldade 2009; Mckinnon and Pierotti 2010). The diversity of insect pigmentation across species, populations, sexes, and individuals of the same sex has been the focus of many eco-evo-devo studies, providing key insight into the genetic basis of variation in pigmentation (e.g. Pool and Aquadro 2007; Futahashi and Fujiwara 2008; Miyagi et al. 2015; Massey and Wittkopp 2016; Zhang et al. 2017; Orteu and Jiggins 2020) and exploring important phenomena such as developmental plasticity (e.g. Solensky and Larkin 2009; Shearer et al. 2016; Monteiro et al. 2020), the origin of novelty (e.g. Shirai et al. 2012; Vargas-Lowman et al. 2019), and evolutionary constraints (Beldade et al. 2002b; Allen et al. 2008).

Variation in body pigmentation can arise from differences in color and/or in the spatial arrangement of colors into specific patterns. These two aspects rely on largely distinct classes of genes involved in pigmentation development: those encoding the enzymes responsible for pigment synthesis, and those encoding the transcription factors regulating those enzymes’ expression at the appropriate time and place (see True 2003; Wittkopp et al. 2003; Wittkopp and Beldade 2009). Changes in genes associated with each of these steps can result in changes in pigmentation between individuals and between body parts (e.g. Wittkopp et al. 2002). In this respect, body pigmentation can be thought of as a multi-dimensional trait, made up of several components representing aspects of actual color and of color pattern, and which might develop and evolve more or less independently. This has been explored in studies focusing on specific color pattern elements, including on butterfly wings (e.g. Nijhout 2001; Monteiro 2015; Beldade and Peralta 2017), as well as on fly wings and abdomens (e.g. Jeong et al. 2006; Werner et al. 2010). Yet, rarely do studies of body pigmentation variation combine quantitative analysis of multiple color and color pattern traits.

Studies of *Drosophila* body and wing pigmentation have provided very valuable insight about the genetic and environmental bases of variation between species, populations of the same species, and individuals of the same population (e.g. Hollocher et al. 2000; Wittkopp et al. 2003; Gibert et al. 2007; Pool and Aquadro 2007; Massey and Wittkopp 2016). These studies characterized effects of environmental factors, such as nutrition (e.g. Shakhmantsir et al. 2014) and temperature (e.g. David et al. 1990), as well as allelic variants of both subtle (e.g. Bastide et al. 2013) and large phenotypic effect (e.g. Carbone et al. 2005). Variation in *Drosophila* pigmentation has been associated to clinal and seasonal variation in desiccation resistance, thermo-regulation, and UV protection (e.g. Rajpurohit et al. 2008; Matute and Harris 2013; Parkash et al. 2014), and shown to correlate with other traits, such as reproductive success, behavior, and immunity (e.g. Dombeck and Jaenike 2004; Takahashi 2013; Massey et al. 2019). While studies of *Drosophila* pigmentation have included focus on different body parts (e.g. trident on thorax, e.g. David et al. 1985; melanic patches on wings, e.g. True et al. 1999; dark bands of abdominal segments, e.g. Dembeck et al. 2015), these studies typically analyze single and often qualitative properties of pigmentation (but see e.g. Saleh Ziabari and Shingleton 2017). Indeed, the detail in quantitative phenotyping of body pigmentation does not match the sophistication of the analysis of its genetic and developmental bases. This is not unique to *Drosophila* pigmentation; the need for more attention to be given to phenotyping has been called for repeatedly (Gerlai 2002; Houle et al. 2010; Kühl and Burghardt 2013; Deans et al. 2015; Laughlin and Messier 2015).

Here, we quantify various traits encompassing aspects of both color and color pattern of abdomen and thorax pigmentation in *Drosophila* adults. We investigate how each of these pigmentation components (or traits) and the associations between them differ between genotypes and developmental temperatures, within and across species. We show that different pigmentation components can vary rather independently, and that fly bodies can be made lighter or darker by changing different pigmentation components. We discuss our results in the context of the potential for evolutionary diversification of pigmentation.

## RESULTS

To investigate patterns and sources of variation in *Drosophila* body pigmentation, we developed a quantitative method to define five pigmentation traits that include aspects of color and color pattern (see Figure S1 and Materials and Methods). We focused on the dorsal surface of thoraxes and abdomens, characterized for having different types of dark “pattern elements” on a lighter “background” color: a trident at the center of the thorax and posterior bands on each segment of the abdomen. Flies were imaged under a binocular scope in controlled light conditions. For each body part, we defined a transect between an anterior and a posterior landmark and collected color information for each pixel along these transects (Figure 1A, Figure S1). Using that information, we quantified a series of traits for each body part: overall darkness (Odk), relative length of transect occupied by the darker “ornamental” pattern (Pat), actual color of both background (Cbk) and “ornamental” pattern elements (Cpa), and the distance in RGB space between the darkest and the lightest that corresponds to the range of color variation (Ran). We investigated how these pigmentation components vary and co-vary between sexes and between rearing temperatures in *D. melanogaster* representing standard laboratory strains, and natural populations from different geographical locations, as well as in five additional *Drosophila* species. For each dataset (*D. melanogaster* laboratory strains, *D. melanogaster* clinal populations, and *Drosophila* species), the multivariate multiple regression analyses showed that pigmentation differed significantly between strains/genotypes/species, sexes, and temperatures, with effects that depended on body part (Table S1).

**Figure 1.**
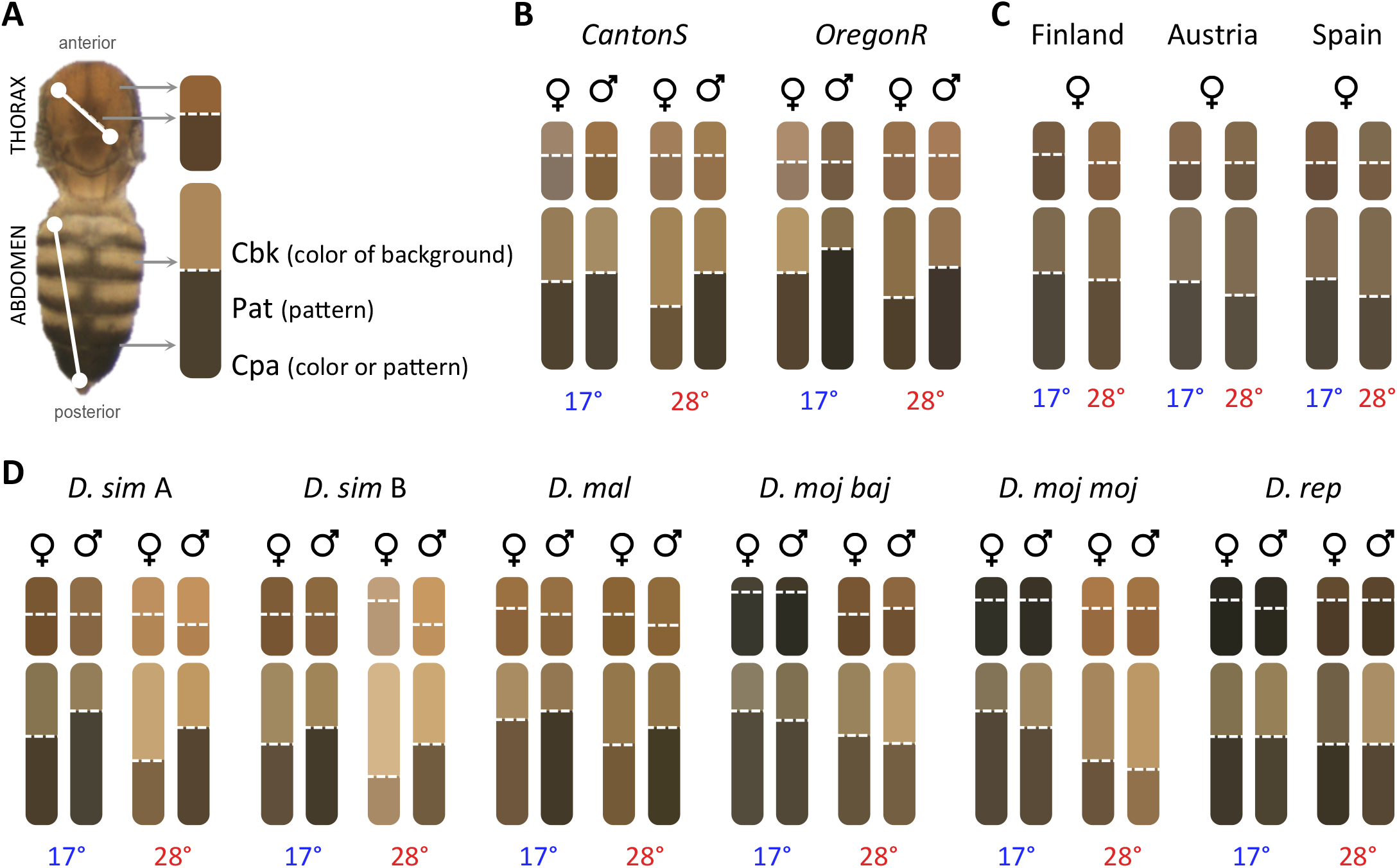
Intra- and inter-specific variation in *Drosophila* pigmentation. **A**. Example of a mounted *D. melanogaster* headless-body showing the dorsal side of the thorax and abdomen with transects, and the scheme we used to represent pigmentation traits for thorax (top rounded rectangle) and abdomen (bottom rounded rectangle). For each of these, the horizontal dashed line separates the color of pattern element (Cpa) and the color of background (Cbk). These are shown in mean color (RGB values) for same-group individuals, and the height of the dashed line represents the proportion of the transect that is occupied by pattern versus background (Pat). See more details in Figure S1. **B**. Pigmentation schemes per strain, sex, and temperature in *D. melanogaster* laboratory populations. **C**. Pigmentation schemes in *D. melanogaster* clinal populations, showing mean values from the five genotypes (i.e. isogenic lines) per location. **D**. Pigmentation schemes in five *Drosophila* species with one genetic background per species except *D. simulans* where two genetic backgrounds (*D. sim* A and *D. sim* B) were studied.

### Variation in body pigmentation in *D. melanogaster* laboratory populations

We reared flies from two common laboratory genetic backgrounds (or strains) of *D. melanogaster*, Oregon R (OreR) and Canton S (CanS), at either 17°C or 28°C to assess thermal plasticity and sexual dimorphism in our pigmentation traits (Figure 1, 2, Figure S2, Table S2). We confirmed known patterns of thermal plasticity and sexual dimorphism for body pigmentation, with flies reared at lower temperature being generally darker than those reared at higher temperature, and males being darker than females (Figure 1B, 2A, Figure S2). However, we found differences between strains and body parts in the extent, and sometimes the direction of both thermal plasticity and sexual dimorphism for our pigmentation traits (Figure 1B, 2A, Figure S2, Table S2), as well as for the correlations between them (Figure 3A).

**Figure 2.**
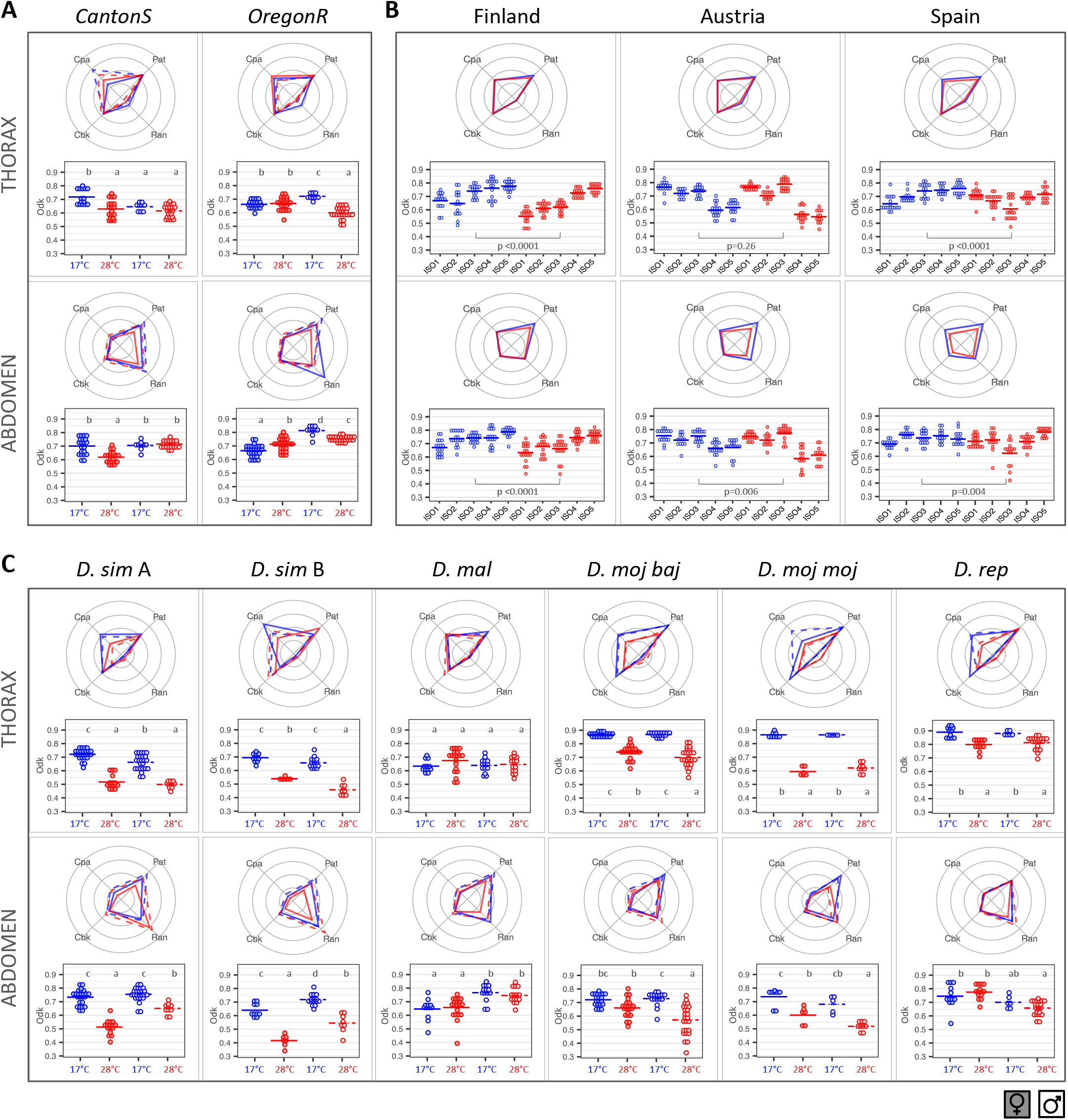
Quantitative phenotyping of *Drosophila* pigmentation component traits. For each population, temperature, sex, and body part, dot plots represent variation for Odk (individual data points and means) and radar plots represent variation for Pat, Ran, Cpa, and Cbk (means; dotplots in Figure S2). Females/males are shown as closed/empty circles (dot plots) or solid/dashed lines (radar plots), and flies reared at 17°C/28°C are shown in blue/red. **A**. *D. melanogaster* laboratory populations. Results of statistical test for the effect of sex, temperature, and their interaction on each of the traits are shown in Table S2. Letters in dot plots indicate results of post-hoc pairwise comparisons between groups: different letters when significantly different (p-value<0.05 for Tukey’s honest significance test). **B**. *D. melanogaster* clinal populations. For each geographical population, we phenotyped females from five genotypes (i.e. isogenic lines). Results for the effect of location, genotype, and temperature (and interactions) on the different pigmentation traits are in Table S4. Results of the statistical test (p-value) for the effect of temperature on each of the traits are shown in plots. **C**. *Drosophila* species. Results of the statistical test for effect of sex, temperature and their interaction are in Table S5. Letters in dot plots indicate results of post-hoc pairwise comparisons between groups: different letters when significantly different (p-value<0.05 for Tukey’s honest significance test).

**Figure 3.**
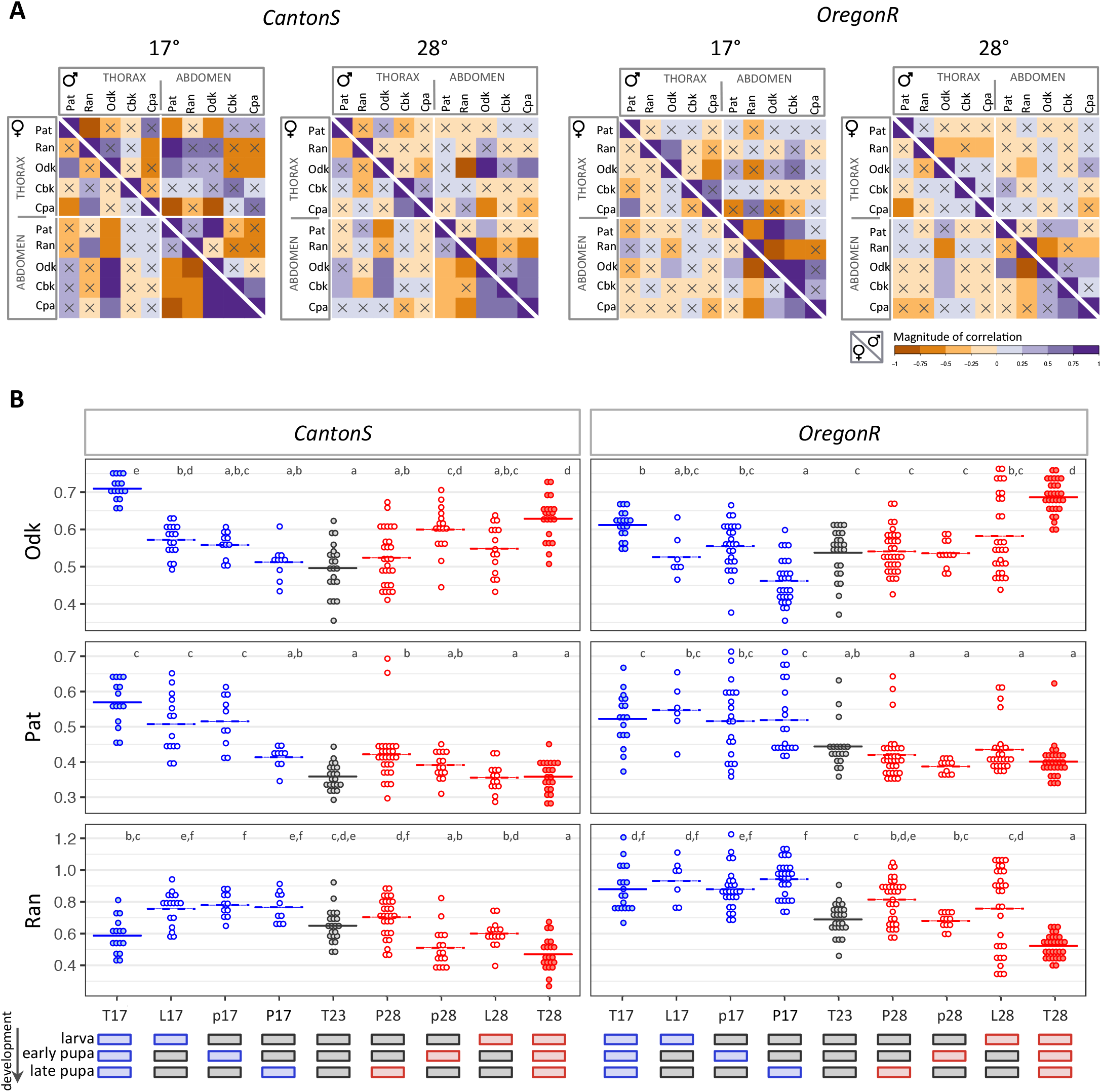
Co-variation and thermal sensitivity of *D. melanogaster* pigmentation components. **A**. Heat map of Pearson’s correlation coefficients for all pigmentation traits in abdomens and thoraxes of CanS (left panels) and OreR (right panels) of flies reared at 17°C or 28°C. For each matrix, females are in the left corner and males in the right. Positive correlations are shown in purple and negative correlations in orange. Correlations not statistically significantly different from zero (p-value > 0.05) are indicated with a cross. **B**. Pigmentation traits (Y axis) in females of two *D. melanogaster* genetic backgrounds (CanS and OreR) exposed to each of the temperature regimes during development (X axis). The thermal regimes codes and corresponding stages that were exposed to either 17°C or 28°C (instead of the basal temperature of 23°C) were: T (constant temperature), L (late larval development), p (early pupal period) and, P (late pupal period). In each graph, dots represent phenotypes of single individual females, and the horizontal bar is the mean of those values. The results of the test for differences between strains and thermal regimes on the different plastic traits are shown in Table S3. Letters indicate results of post-hoc pairwise comparisons between groups: different letters when significantly different (p-value<0.05 for Tukey’s honest significance test).

For overall darkness (Odk; dot plots in Figure 2A), flies reared at 17°C were generally darker than those from 28°C, with the exception of CanS males (where differences were not significant in either body part), and OreR females (where abdomens were darker in flies from 28°C). The abdomens were lighter in females relative to males (except for CanS from 17°C), but the thoraxes were lighter in males relative to females (except for CanS from 28°C and OreR from 17°C). We also observed differences between sexes and temperatures for the other pigmentation traits (Pat, Ran, Cbk, and Cpa; radar plots in Figure 2A; dot plots in Figure S2, Table S2), which depended on body part. While for the thorax the most striking differences were seen in Ran (for females between temperatures), for the abdomen they were seen for Pat (distinguishing females from 28°C from others) and Ran (extreme for OreR females) (Figure S2). Variation was only loosely correlated between traits, with few significant correlations, which differed between genetic backgrounds, sexes, and rearing temperatures. Overall, correlations between traits were weaker across body parts relative to within body parts, and in males relative to females (Figure 3A).

For those pigmentation traits found to be thermally plastic (i.e. significant differences between individuals reared at different temperatures; cf. Figure S2, Table S2), we investigated which stages of development were thermally responsive. To do so, we compared phenotypes between individuals (specifically, female abdomens) differing in temperature only for specific developmental time windows (Figure 3B, Figure S3, Table S3). We tested nine thermal regimes (i.e. treatments), including three with constant temperatures (whole development at 17°C, 23°C, or 28°C) and six where most of the development took place at 23°C and only one specific stage (either late larval, early pupal, or late pupal) took place at 17°C or at 28°C. Differences between constant temperatures (T17, T23, and T28 treatments), revealed thermal reaction norms, i.e. the representation of phenotype as a function of temperature (see Schlichting and Pigliucci 1998), of different shapes for different pigmentation components: T23 phenotype intermediate between T17 and T28 (Ran in OreR; Figure 3B), equal to one of the extreme temperatures (Pat; Figure 3B), or more extreme than both T17 and T28 (Odk; Figure 3B). The period when exposure to a different temperature significantly affected phenotype also differed between traits and genetic backgrounds (Figure 3B, Figure S3).

### Body pigmentation differences between *D. melanogaster* natural populations and *Drosophila* species

We quantified variation in pigmentation components in wild-caught populations sampled along a latitudinal cline in Europe: Finland, Austria, and Spain (samples from the *DrosEU Consortium;* http://droseu.net/). We analyzed pigmentation traits in females from five genotypes (isofemale lines) established from each of the three geographical locations, reared at either 17°C or 28°C. The analysis for each pigmentation component revealed differences between traits in their response to the various explanatory variables and their interactions (Figure 1C, 2B, Table S4). Geographical populations differed in overall darkness (Odk; dot plots in Figure 2B) and in color (both Cbk and Cpa) for the abdomen, but not the thorax (Figure 2B, Figure S5, Table S4). For the thorax, only Ran and Pat differed between locations (Figure 2B, Table S4). Most pigmentation traits (except thorax color; Cpa and Cbk) were thermally plastic, with darker flies for development at 17°C relative to 28°C (Figure 1C, 2B, Figure S4). The Northern- and Southern-most populations (i.e. Finland and Spain, respectively) did not necessarily show the most extreme phenotypes, neither in terms of overall darkness nor in the extent of plasticity therein (Figure 2B, Figure S5). We also found significant differences between isofemale genotypes (and their plasticity) within each geographical location (Figure 2B, Table S4).

Finally, we quantified pigmentation traits in flies from five additional *Drosophila* species (two genetic backgrounds for *D. simulans*, and one genetic background for all other species or sub-species: *D. malerkotliana, D. repleta*, *D. mojavensis baja*, *D. mojavensis mojavensis*) reared at either 17°C or 28°C (Figure 1D, 2C). We found differences between species in extent and direction of sexual dimorphism and of thermal plasticity for the different pigmentation traits (Figure 1D, 2C, Figure S2B, Table S5). For instance, for Odk (dot plots in Figure 2C), while *D. malerkotliana* showed no differences between temperatures and clear differences between sexes, *D. simulans* had very high thermal plasticity but reduced sexual dimorphism (no differences between females and males reared at 17°C). For the other pigmentation traits (radar plots in Figure 2C and dot plots in Figure S2B), larger differences between sexes and/or temperatures were observed for Pat and/or Ran, and less for actual colors (Cpa and Cbk).

## DISCUSSION

We decomposed *Drosophila* body pigmentation into different quantitative traits, including overall darkness (Odk), and traits reflecting properties of color and color pattern (Pat, Ran, Cbk, and Cpa) of both thoraxes and abdomens. We showed differences in trait values, as well as in the extent and direction of thermal plasticity and of sexual dimorphism for laboratory and natural populations of *D. melanogaster* and across *Drosophila* species (Figures 1, 2). Different traits, corresponding to different properties of body pigmentation, behaved in a largely independent manner, which was also reflected in low levels of correlations between traits and in differences in the period of development during which traits are thermally responsive (Figure 3).

*Drosophila* pigmentation has been the focus of various studies exploring aspects of its ecology, development and evolution (e.g. Kopp et al. 2000; Williams et al. 2008; Matute and Harris 2013; Shearer et al. 2016; Gibert et al. 2017). This has provided great insight about the genetic basis and ecological significance of variation, across temporally (e.g. seasonal variation) or geographically (e.g. clinal variation) distinct populations (e.g. Parkash et al. n.d.; Hollocher et al. 2000b; Rajpurohit et al. 2008), as well as across species (Hollocher et al. 2000a,b). Many of those studies focused on specific pigmentation elements in particular species, and often used qualitative assessments of pigmentation variation or presence/absence of specific pattern elements (e.g. Hollocher et al. 2000; David et al. 2002). In *D. melanogaster* for instance, most work has focused on abdominal pigmentation, and specifically on the dark bands of the posterior-most segments, which is sexually dimorphic (males are generally darker than females; e.g. Kopp et al. 2000) and thermally plastic (flies from lower developmental temperatures are generally darker than flies from higher developmental temperatures; e.g. David et al. 1990; Gibert et al. 2007, 2009). We extended the analysis of body pigmentation to quantifying different properties of both abdomen and thorax pigmentation in *D. melanogaster* and other *Drosophila* species. This more detailed analysis ultimately painted a more complex picture of variation in *Drosophila* body pigmentation. We did not, for instance, always find that males were darker than females, or that flies reared at lower temperatures were darker than those from higher temperatures, but rather, we found trait specificities in how pigmentation varied between sexes and between developmental temperatures. This was true for overall darkness (Odk) of the abdomen, the trait that would presumably be more similar to previous (largely qualitative) characterizations of abdominal pigmentation (e.g. David et al. 1990; Hollocher et al. 2000a), but also for other properties of body pigmentation, including actual color of background and pattern elements (i.e. abdominal bands and thoracic trident). Moreover, we also showed that pigmentation components, as well as sexual dimorphism and thermal plasticity therein, vary greatly between species, genotypes, and body parts. The mechanisms underlying such intra- and inter-specific variation in different traits, as well as the trait-specific responses to temperature, remain to be explored and might involve differences in the environmental sensitivities of the regulatory regions (e.g. enhancers) controlling pigmentation-related genes (e.g. De Castro et al. 2018).

Our results also highlight that the different components vary largely independently, with only weak correlations between traits (Figure 3A) and differences between traits in the extent and direction of thermal plasticity and of sexual dimorphism (Figure 1, 2). Pigmentation components were shown to even differ in the period of development in which they are responsive to temperature (Figure 3B). Similar environmental effects on trait associations have been described previously; for instance, cold temperature triggered a shift in the sign of the correlation between body size and longevity in *D. melanogaster* (Norry and Loeschcke 2002). Differing correlations between body parts (or regions within a body part) have also been identified for *D. melanogaster* pigmentation (e.g. Gibert et al. 2000; Bastide et al. 2014), with the extent of genetic correlations decreasing with increasing distance between body segments (Gibert et al. 2000). Ultimately, the dependency of trait associations on genetic and environmental factors has the potential to influence adaptation (e.g. Marquez & Knowles 2007; Manenti *et al*. 2016), as evolutionary change can result from both direct and correlated responses to selection (e.g. Rajpurohit and Gibbs 2012). Altogether, our results suggest a large degree of developmental and evolutionary independence between pigmentation components, which could facilitate the diversification of body coloration in *Drosophila*.

Studies exploring the ecological conditions driving the evolution of melanism in *Drosophila* have documented correlations between body pigmentation and several eco-geographic variables (e.g. latitude, altitude, temperature, humidity) (e.g. Rajpurohit et al. 2008; Gibert et al. 2016; Shearer et al. 2016). Clinal variation in pigmentation, for instance, has been shown for thoracic trident (e.g. David et al. 1985; Telonis-Scott et al. 2011) and for abdominal pigmentation (e.g. Pool and Aquadro 2007; Das 2009). Generally, darker phenotypes in colder environments (e.g. at high latitudes or altitudes) have been hypothesized to allow flies to better absorb solar radiation (c.f. thermal budget or thermal melanism hypothesis; Trullas et al. 2007; Clusella-Trullas et al. 2008), to increase desiccation resistance (e.g. Parkash et al. 2008), and/or to provide protection against UV radiation (e.g. Bastide et al. 2014). Plasticity, on the other hand, is expected to be greater in environments that are more variable (Lande 2014), such as those with larger seasonal fluctuations, often occurring at higher latitudes. However, our analysis of the pigmentation patterns from *D. melanogaster* populations collected along a European latitude cline (Finland, Austria, Spain) did not always revealed darker pigmentation nor higher plasticity in the Northern-most population (i.e. Finland), which may indicate that other environmental parameters and ecological conditions not considered here could account for the differences between populations in the different pigmentation components. Having only three populations from three latitudes may also be limiting in terms of assessing latitudinal patterns in pigmentation and plasticity therein.

In terms of a function in thermo-regulation favoring darker flies in cooler environments (David et al. 1985; Hollocher et al. 2000a; Wittkopp et al. 2011; Matute and Harris 2013; Shearer et al. 2016), we could expect our trait overall darkness (Odk) to be the most relevant trait. Our analyses revealed that flies can become overall darker (higher Odk) by changing actual colors of background or of pattern elements (Cbk and Cpa, respectively) or the proportion of the abdomen/thorax length covered with the darker bands/trident (Pat). For instance, males of CanS reared at 17°C and 28°C, show the same overall darkness (Odk), but differ in what pigmentation components make that up; Odk is mostly determined by color components at 17°C and by color pattern components at 28°C (i.e. Cpa and Cbk are lower, while Pat and Ran are higher at 17°C than at 28°C). It is unclear whether these traits are mere components of Odk or are themselves under direct natural selection.

Variation in pigmentation between body parts, individuals, populations, and species can be caused by differences in actual color and/or in how colors are spatially organized to make up color patterns (Wittkopp and Beldade 2009; Nijhout 2010). However, seldom do studies of animal pigmentation consider and quantify distinct pigmentation component traits, and the extent to which they might be differently affected by genetic and/or environmental variation. The increased attention to studying the mechanisms underlying phenotypic variation resulted in great detail and sophistication in the characterization of its genetic underpinnings. However, the detail in describing and quantifying phenotypes has lagged behind. The lack of quantitative methods for phenotyping (see Gerlai 2002; Houle et al. 2010) can result in an oversimplification of complex phenotypes, dismissing that those phenotypes are often made up of distinct component traits that can respond to internal and external factors in different manners (e.g. Vrieling et al. 1994; Mateus et al. 2014). We attempted to provide a better resolution of variation in *Drosophila* body color, a visually compelling example of adaptive evolution. Combining it with existing genetic resources and with access to natural variation can provide a deeper resolution of the patterns and processes underlying phenotypic variation, within and between species.

## MATERIAL AND METHODS

### Fly stocks

*D. melanogaster* genetic backgrounds CantonS (CanS) and OregonR (OreR) and *Drosophila* species *D. simulans, D. malerkotliana, D. repleta*, *D. mojavensis baja* and *D. mojavensis mojavensis* were obtained from C. Mirth’s lab. *D. melanogaster* populations from Finland (Akaa; 61.1, 23.52; collected in July 2015), Austria (Mauternbach; 48.38, 15.57; collected in July 2016) and Spain (Tomelloso; 39.16, 3.02; collected in September 2015) were obtained from E. Sucena’s lab and collected by members of the *European Drosophila Population Genomics Consortium* (*DrosEu*; http://droseu.net). All stocks were maintained in molasses food (45 gr. molasses, 75 gr. sugar, 70 gr. cornmeal, 20 gr. Yeast extract, 10 gr. Agar, 1100 ml water and 25 ml of Niapagin 10%). All stocks were kept at 25°C, 12:12 light-dark cycles. For the experiments, we performed over-night egg-laying from ~20 females of each stock in vials with *ad libitum* molasses food. Eggs were then placed at either 17°C or 28°C throughout development. We controlled the population density by keeping between 20 and 40 eggs per vial.

For the experiment of the windows of sensitivity for pigmentation, we exposed developing flies to 17°C or 28°C during one window of development while they were kept at 23°C for the remaining stages. We tested four different treatments at 17°C and at 28°C: T (flies always kept at constant temperature), L (late larval development; staging done by using traqueal and mouth hook morphology), p (only early pupal period; from white pupa to the onset of eye pigmentation), P (only late pupal period; from the onset of eye pigmentation until adult eclosion).

### Phenotyping pigmentation components

Adult flies (8-10 days after eclosion) were placed in 2 ml microcentrifuge tubes and frozen in liquid nitrogen. The tubes were shaken immediately after submersion in liquid nitrogen to remove wings, legs and bristles. Headless bodies of flies were then mounted on 3% Agarose in Petri dishes, dorsal side up, and covered with water to avoid specular reflection of light upon imaging. Images containing 10 to 20 flies were collected with a LeicaDMLB2 stereoscope and a Nikon E400 camera under controlled conditions of illumination and white-balance adjustment. Images were later processed with a set of custom-made interactive Mathematica notebooks (Wolfram Research, Inc., Mathematica, Version 10.2, Champaign, IL, 2015) to extract pigmentation measurements. For this purpose, two transects were defined on each fly, one in the thorax and one in the abdomen, using morphological landmarks (as shown in Figure S1). To minimize image noise, for each pixel position along the transect line we calculated the mean RGB (Red, Green, Blue) values of the closest five pixels located on a small perpendicular line centered on the transect. For abdominal transects, when necessary, we removed the sections corresponding to the membranous tissue that occasionally is visible between abdominal segments. The few transects that were drawn over debris particles were excluded from the analysis, as pigmentation measurements could not be accurately extracted.

The sequence of averaged RGB pixel values corresponding to each transect was then used to define each of the five pigmentation components as follows. For each pixel, we calculated a normalized darkness value as Dmax–Dbk, where Dmax is the largest possible Euclidean distance between two colors in the RGB color space (in this case 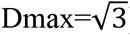), and Dbk is the distance of the pixel’s color coordinates to the color black (R=0, G=0, B=0). Overall darkness (Odk) was calculated as the sum of the normalized darkness values for each pixel divided by the number of pixels in the transect. Taking the sequence of normalized darkness values along a transect, we estimated its two enveloping lines (blue and green lines in Figure S1A) by calculating the baselines of the original and negated values using the Statistics-sensitive Non-linear Iterative Peak-clipping (SNIP) algorithm (Ryan et al. 1988). The median line of this envelope (red line in Figure S1A) was then used to separate the transect pixels into two clusters, where the pixels above or below this line correspond, respectively, to the pattern element (trident in the thorax and darker bands in the abdomen) or to the background. Pattern (Pat) was calculated as the proportion of pixels corresponding to the pattern element relative to the transect length. Color of the pattern element (Cpa) is the angle defined in the RGB color space between the best-fitted line going through the color coordinates of the pixels in the transect that correspond to the pattern element (trident and/or darker bands) in the transect and the gray vector (the black to white diagonal in the RGB color space). Similarly, color of the background (Cbk) was calculated as the angle between the best-fitted line that goes through the color coordinates of the background pixels in the transect and the gray vector. Pixels corresponding to pattern element and/or background were defined by grouping all RGB values in the transect into two clusters each containing 95% of the light or dark pixels respectively. Range (Ran) was calculated as the Euclidean distance between the median values of the 20 darkest and the 20 lightest pixels along the transects. The colors represented in Figure 1 correspond to the mean R, mean G and mean B values for each strain/species, sex, and temperature, which were calculated from Cpa for color of pattern elements and from Cbk for color of the background, respectively.

### Statistical analyses

All analyses were conducted in R v 3.6.2 (R Core Team 2019), using the following R packages: *tidyr* (Wickham and Henry 2020) to arrange datasets, *ggplot2* (Wickham 2009) to produce all plots, *lme4* (Bates et al. 2015) and *lmerTest* (Kuznetsova et al. 2017) to perform linear mixed-effects models, *corrplot* (Taiyun and Viliam 2017) to compute correlation matrices, and *emmeans* (Lenth et al. 2018) to perform post-hoc pairwise comparisons between groups. The statistical models described below are given in package-specific R syntax (shown in italics).

Multivariate multiple regression was performed for the data on *D. melanogaster* laboratory populations to test for the effect of strain, sex, temperature (fixed explanatory variables), and interaction terms in all pigmentation traits by combining all traits using the *cbind* function (model lm*(cbind(Odk, Pat, Ran, Cbk, Cpa) ~ Strain * Sex * Temperature))*. A similar analysis was performed for the data on *D. melanogaster* clinal populations testing for the fixed effects and interactions of location, genotype (i.e. isogenic line; nested within location), and temperature (model *lm(cbind(Odk, Pat, Ran, Cbk, Cpa) ~ Location * Genotype * Temperature*), and for the *Drosophila* species, testing for the fixed effects and interactions of species, strain (nested within species), sex, and temperature (model lm*(cbind(Odk, Pat, Ran, Cbk, Cpa) ~ Species * Species/Strain * Sex * Temperature))*, where *Strain* corresponds to the different genetic backgrounds analyzed in *D. melanogaster* (CanS and OreR) and in *D. simulans* (*D. sim* A and *D. sim* B).

Linear mixed effect models were then used to test for the (fixed) effects of different explanatory fixed variables (strains, genotypes or species, sex and temperature) and their interactions on each of the pigmentation traits (noted as *trait* in the model notations below). *Replicate* was included as random effect in the models (corresponding to the *(1|Replicate)* factor in the R syntax below). For *D. melanogaster* laboratory strains: model lm*(Trait ~ Sex * Temperature + (1|Replicate)))*. The same model was used for all *Drosophila* species, except for *D. simulans*, where we also included the factor *Strain* which corresponds to the different genetic backgrounds studied in this species (*D. sim* A and *D. sim* B) (model: lm(*Trait ~ Strain * Sex * Temperature + (1|Replicate)*)). For the clinal populations: model: lm*(Trait ~ Location * Location/Genotype * Temperature + (1|Replicate))*. For all the aforementioned mixed models, we used Satterthwaite's method (via *anova* function in lmerTest package; Kuznetsova et al. 2017) for approximating degrees of freedom and estimating F-statistics and P-values. For the data on the sensitive stages of development, we used linear effect models to test for the effect and interaction of strain and thermal regime (model: lm*(Trait ~ Strain*Regime))*.

We used *post-hoc* pairwise comparisons (Tukey’s honest significant differences) to identify differences between strains, sexes, temperatures and/or thermal regimes. Pearson’s correlations were used to check correlations between traits and across temperatures.

## DATA ACCESIBILITY

All data will be made publicly available in Dryad Digital Repository upon acceptance of the manuscript.

## AUTHOR CONTRIBUTIONS

E.L. and P.B. conceived and designed the study. E.L., J.G.K., and C.M.P. performed the experiments. F.A. developed the quantitative method for color pattern analysis and the respective computational tools. E.L. analyzed the data. E.L. and P.B. wrote the manuscript.

## ACKNOWLEDGMENTS

We are grateful to Emanuel Fernandes for collecting preliminary data on the developmental windows of thermal sensitivity, to Christen Mirth for sharing the *Drosophila* species, to Élio Sucena and the *Drosophila Population Genomics Consortium* (*DrosEU*; http://droseu.net/), funded by a Special Topic Networks (STN) grant from the European Society for Evolutionary Biology (ESEB), for access to the *D. melanogaster* European populations. We thank Gabriel Martins and the Imaging Facility at the IGC for support in setting up the image acquisition system.

