## Supplementary Material for "Many ways to make darker flies: Intra- and inter-specific variation in *Drosophila* body pigmentation components"

### SUPPLEMENTARY FIGURES

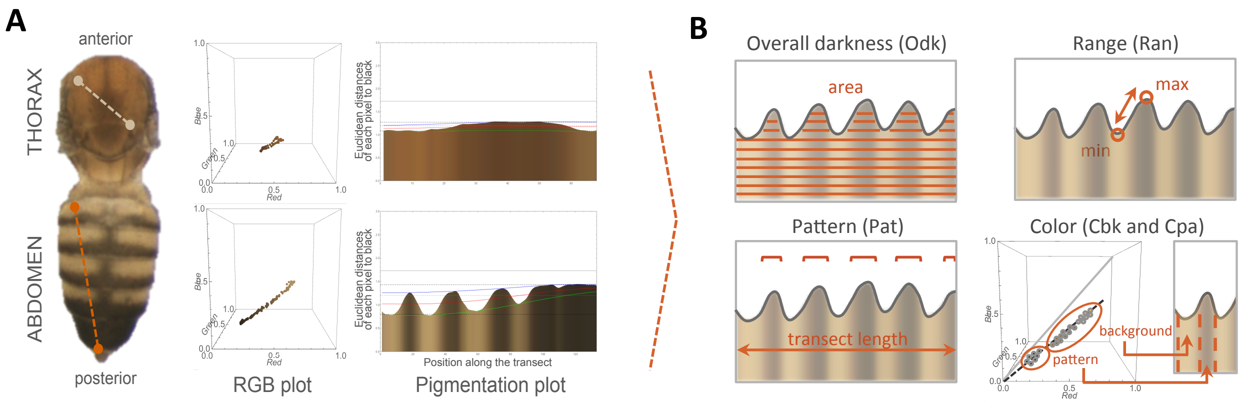


**Figure S1. Quantitative analysis of body pigmentation. A.** Thorax and abdomen from female of *D. melanogaster* OreR reared at 17°C showing the body landmarks used to draw the transects. For each pixel in the transect we extracted RGB values that are represented in the RGB plots (cubes on the right side of each transect). By calculating the distance between each of those pixels to the black, we converted the RGB vectors into two dimensional information and represented the distance of each pixel (Y axis) from the anterior to the posterior extremes of the transect (X axis) (plots on the right side). **B.** Diagram showing the different pigmentation traits. Overall darkness (Odk), difference between lightest and darkest color (Ran), relative length of dark “ornamental” pattern (Pat), color of background (Cbk), and color of pattern “ornamental” elements (Cpa) (see Materials and Methods).

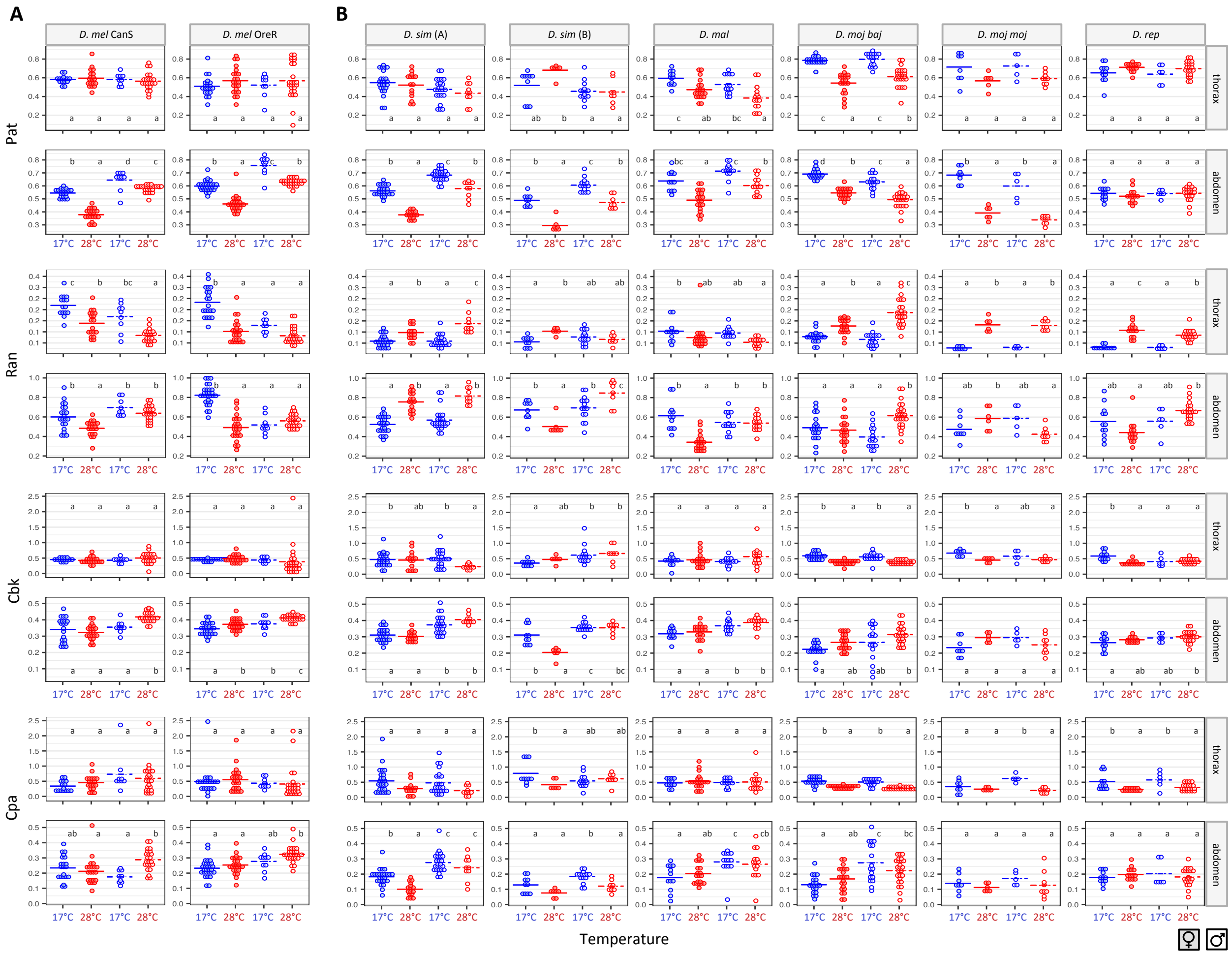


**Figure S2. Variation in pigmentation traits with sex and temperature in *Drosophila*.** For each population/species, temperature, sex, and body part, dot plots represent variation for pigmentation traits Pat, Ran, Cbk, and Cpa (individual data points and means, represented with bar). Females/males are shown as closed/empty circles and flies reared at 17°C/28°C are shown in blue/red. **A.** *D. melanogaster* laboratory populations. Results of statistical test for the effect of sex, temperature, and their interaction on each of the traits are shown in Table S2. Letters in dot plots indicate results of post-hoc pairwise comparisons between groups: different letters when significantly different (p-value<0.05 for Tukey's honest significance test). **B.** *Drosophila* species. Results of the statistical test for effect of sex, temperature and their interaction are in Table S5. Letters in dot plots indicate results of post-hoc pairwise comparisons between groups: different letters when significantly different (p-value<0.05 for Tukey's honest significance test). For *D. simulans* (*D. sim*), we had two different strains (A and B), which are shown independently in the graph and were included in the statistical model.

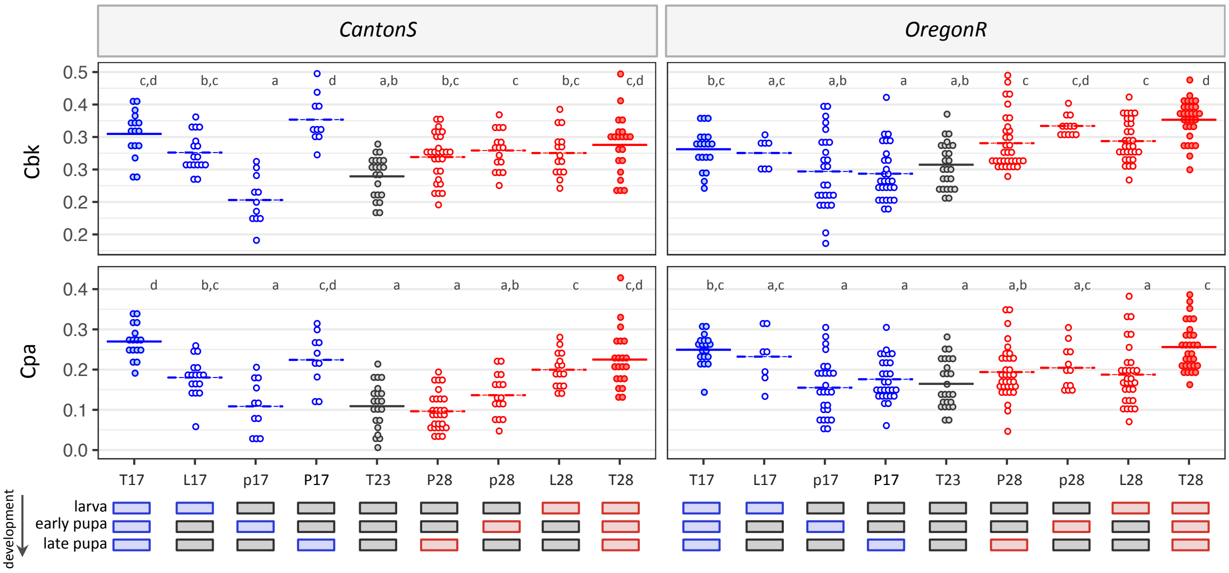


**Figure S3. Windows of sensitivity for pigmentation plasticity in *D. melanogaster*.** Pigmentation traits Cbk and Cpa (Y axis) in females of two *D. melanogaster* genetic backgrounds (CanS and OreR) exposed to each of the thermal regimes (X axis). The codes for the thermal regimes and corresponding stages that were exposed to either 17°C or 28°C degrees were: T (constant temperature), L (late larval development), p (early pupal period) and, P (late pupal period). In each graph, dot plots represent variation for pigmentation traits (individual data points and means, represented with bar). Treatments at 17°C/28°C are shown in blue/red. Results of statistical test for the effect of strain and thermal regime (i.e. treatment) are shown in Table S3. Letters in dot plots indicate results of post-hoc pairwise comparisons between treatments: different letters when significantly different (p-value<0.05 for Tukey's honest significance test).

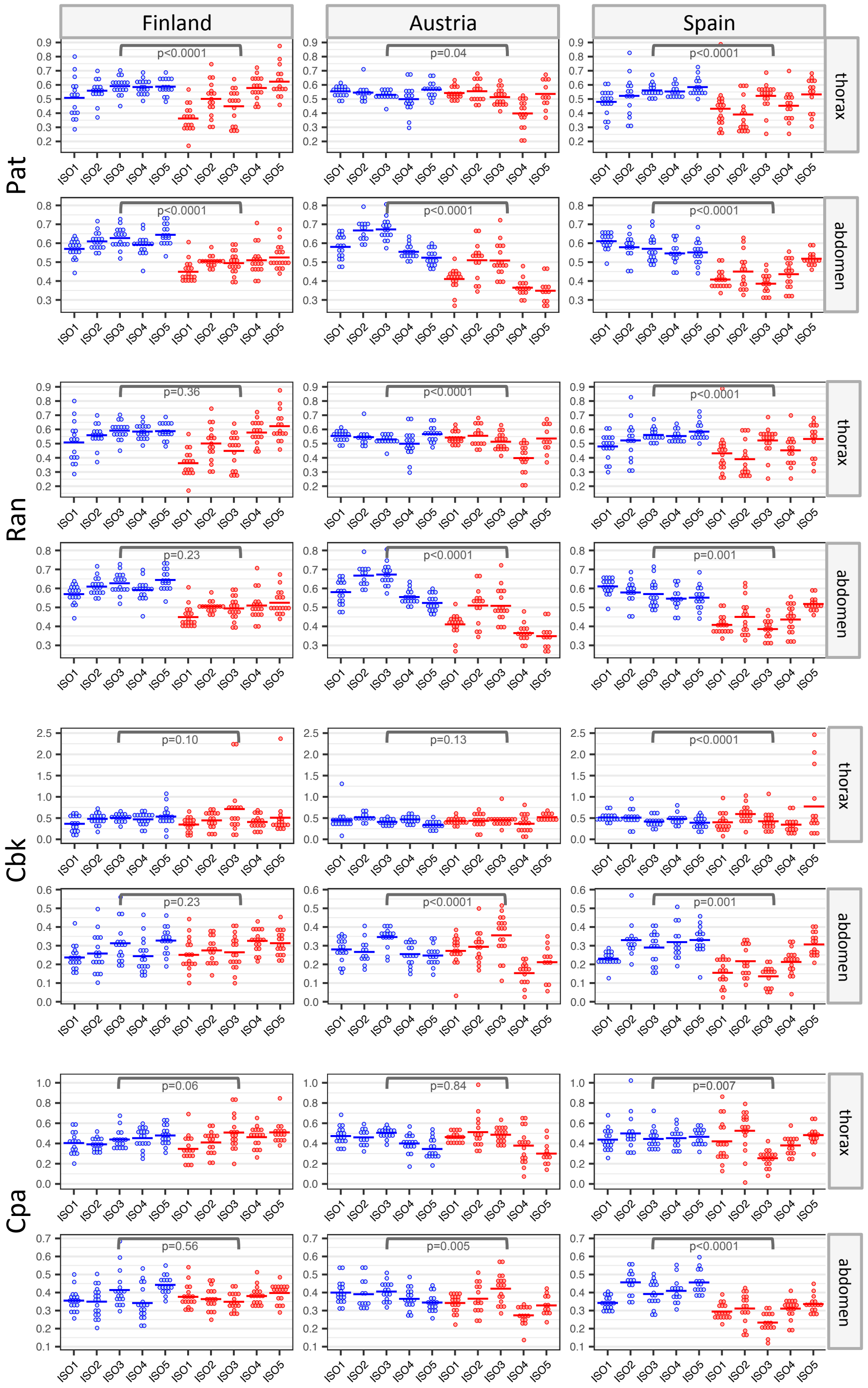


**Figure S4. Variation in pigmentation traits in *D. melanogaster* European populations.** In each graph, dot plots represent variation for pigmentation traits (individual data points and means, represented with bar) with flies reared at 17°C/28°C are shown in blue/red. For each geographical population we phenotyped females from five genotypes (i.e. isogenic lines). Results of statistical test for the effect of location, genotype and temperature are shown in Table S4. Results of the statistical test (p-value) for the effect of temperature on each of the traits are shown in plots.

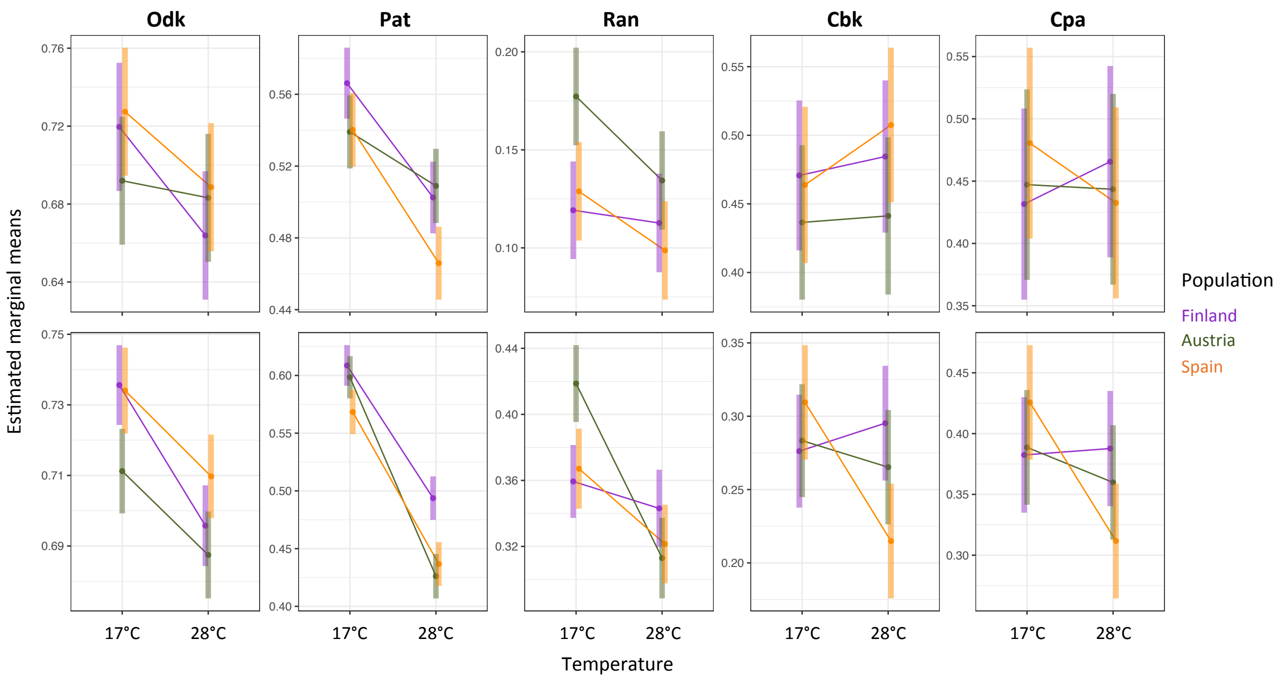


**Figure S5. Effects of temperature on pigmentation traits in *D. melanogaster* European populations.** Interaction plot showing the estimated marginal means and confidence intervals of all pigmentation traits based on fitted model lmer*(Trait ~ Location * Location/Genotype * Temperature + (1|Replicate))*.

### SUPPLEMENTARY TABLES

**Table S1. Pigmentation in *D. melanogaster* populations and *Drosophila* species.** Results of analysis of variance for multivariate multiple regressions performed on each dataset, for the effects of different factors on pigmentation of each body part. Non-significant effects (p-value > 0.05) are shown in grey. For *D. melanogaster* laboratory populations, table shows the results for the effect and interactions of strain, sex, and temperature (model: *lm(cbind(Pat, Ran, Odk, Cbk, Cpa) ~ Strain * Sex * Temperature)*) on pigmentation of each body part. For *D. melanogaster* European populations, table shows the results for the effect and interactions of location, genotype (i.e. isogenic line), and temperature (model *lm(cbind(Odk, Pat, Ran, Cbk, Cpa) ~ Location * Genotype * Temperature)*). For *Drosophila* species, table shows the results for the effect and interactions of species, strain (nested within species), sex, and temperature (model *lm(cbind(Odk, Pat, Ran, Cbk, Cpa) ~ Species * Species/Strain * Sex * Temperature))*. The factor *Strain* corresponds to the different genetic backgrounds analyzed in *D. melanogaster* (CanS and OreR) and in *D. simulans* (*D.sim* A and *D.sim* B).

| | | THORAX | | | | | | ABDOMEN | | | | | |
|---|---|---|---|---|---|---|---|---|---|---|---|---|---|
| | | Df | Pillai | approx F | num Df | den Df | Pr(>F) | Df | Pillai | approx F | num Df | den Df | Pr(>F) |
| D. melanogaster laboratory populations | (Intercept) | 1 | 1,00 | 7342,50 | 5 | 132 | < 2,2E-16 | 1 | 1,00 | 41547,00 | 5 | 153 | < 2,2E-16 |
| | Strain | 1 | 0,04 | 1,10 | 5 | 132 | 3,63E-01 | 1 | 0,58 | 42,00 | 5 | 153 | < 2,2E-16 |
| | Temperature | 1 | 0,65 | 49,70 | 5 | 132 | < 2,2E-16 | 1 | 0,69 | 69,00 | 5 | 153 | < 2,2E-16 |
| | Sex | 1 | 0,33 | 12,70 | 5 | 132 | 4,11E-07 | 1 | 0,84 | 162,00 | 5 | 153 | < 2,2E-16 |
| | Strain:Temperature | 1 | 0,03 | 0,80 | 5 | 132 | 5,42E-01 | 1 | 0,22 | 9,00 | 5 | 153 | 2,20E-04 |
| | Strain:Sex | 1 | 0,05 | 1,40 | 5 | 132 | 2,17E-01 | 1 | 0,38 | 19,00 | 5 | 153 | 1,48E-11 |
| | Temperature:Sex | 1 | 0,08 | 2,20 | 5 | 132 | 6,19E-02 | 1 | 0,55 | 38,00 | 5 | 153 | < 2,2E-16 |
| | Strain:Temperature:Sex | 1 | 0,20 | 6,70 | 5 | 132 | 1,24E-02 | 1 | 0,26 | 11,00 | 5 | 153 | 7,47E-06 |
| | Residuals | 136 | | | | | | 157 | | | | | |
| D. melanogaster European populations | (Intercept) | 1 | 1,00 | 26858,70 | 5 | 456 | < 2,2E-16 | 1 | 9,98E-01 | 59146 | 5,00 | 464 | < 2,2E-16 |
| | Location | 2 | 0,22 | 11,10 | 10 | 914 | < 2,2E-16 | 2 | 2,39E-01 | 13 | 10,00 | 930 | < 2,2E-16 |
| | Temperature | 1 | 0,28 | 35,10 | 5 | 456 | < 2,2E-16 | 1 | 6,76E-01 | 193 | 5,00 | 464 | < 2,2E-16 |
| | Location:Genotype | 12 | 1,15 | 11,40 | 60 | 2300 | < 2,2E-16 | 12 | 9,56E-01 | 9 | 60,00 | 2340 | < 2,2E-16 |
| | Location:Temperature | 2 | 0,11 | 5,50 | 10 | 914 | 4,99E-08 | 2 | 3,02E-01 | 17 | 10,00 | 930 | < 2,2E-16 |
| | Location:Genotype:Temperature | 12 | 0,45 | 3,80 | 60 | 2300 | < 2,2E-16 | 12 | 4,20E-01 | 4 | 60,00 | 2340 | < 2,2E-16 |
| | Residuals | 460 | | | | | | 468 | | | | | |
| Drosophila species | (Intercept) | 1 | 1,00 | 29882,00 | 5 | 450 | <2,2e-16 | 1 | 1,00 | 62309,00 | 5 | 471 | <2,2e-16 |
| | Species | 5 | 1,19 | 28,40 | 25 | 2270 | <2,2e-16 | 5 | 1,26 | 32,00 | 25 | 2375 | <2,2e-16 |
| | Temperature | 1 | 0,60 | 136,30 | 5 | 450 | <2,2e-16 | 1 | 0,69 | 213,00 | 5 | 471 | <2,2e-16 |
| | Sex | 1 | 0,13 | 13,70 | 5 | 450 | 1,80E-09 | 1 | 0,63 | 158,00 | 5 | 471 | <2,2e-16 |
| | Species:Strain | 2 | 0,05 | 2,50 | 10 | 902 | 6,87E-03 | 2 | 0,34 | 19,00 | 10 | 944 | <2,2e-16 |
| | Species:Temperature | 5 | 0,95 | 21,30 | 25 | 2270 | <2,2e-16 | 5 | 0,64 | 14,00 | 25 | 2375 | <2,2e-16 |
| | Species:Sex | 5 | 0,25 | 4,70 | 25 | 2270 | 1,07E-10 | 5 | 0,76 | 17,00 | 25 | 2375 | <2,2e-16 |
| | Temperature:Sex | 1 | 0,02 | 2,30 | 5 | 450 | 4,44E-02 | 1 | 0,27 | 35,00 | 5 | 471 | <2,2e-16 |
| | Species:Strain:Temperature | 2 | 0,05 | 2,20 | 10 | 902 | 1,82E-02 | 2 | 0,24 | 13,00 | 10 | 944 | <2,2e-16 |
| | Species:Strain:Sex | 2 | 0,07 | 3,00 | 10 | 902 | 8,57E-04 | 2 | 0,18 | 10,00 | 10 | 944 | 2,62E-12 |
| | Species:Temperature:Sex | 5 | 0,11 | 2,00 | 25 | 2270 | 2,66E-03 | 5 | 0,30 | 6,00 | 25 | 2375 | <2,2e-16 |
| | Species:Strain:Temperature:Sex | 2 | 0,12 | 5,70 | 10 | 902 | 2,33E-05 | 2 | 0,10 | 5,00 | 10 | 944 | 4,21E-04 |
| | Residuals | 454 | | | | | | 475 | | | | | |

**Table S2. Pigmentation components in *D. melanogaster* laboratory populations**. Results of analysis of variance for the effect and interaction of sex and temperature on the different pigmentation traits per body part (model: *lm(Trait ~ Sex * Temperature + (1|Replicate)*). Non-significant effects (p-value > 0.05) are shown in grey.

| | | | THORAX | | | | | | ABDOMEN | | | | | |
|---|---|---|---|---|---|---|---|---|---|---|---|---|---|---|
| | | | **Sum Sq** | **Mean Sq** | **NumDF** | **DenDF** | **F value** | **Pr(>F)** | **Sum Sq** | **Mean Sq** | **NumDF** | **DenDF** | **F value** | **Pr(>F)** |
| **CantonS** | **Pat** | **Sex** | 0,00 | 0,00 | 1 | 60,98 | 0,08 | 7,84E-01 | 0,29 | 0,29 | 1 | 69,49 | 186,01 | <2,2E-16 |
| | | **Temperature** | 0,01 | 0,01 | 1 | 60,84 | 0,99 | 3,24E-01 | 0,16 | 0,16 | 1 | 69,02 | 101,82 | 3,20E-15 |
| | | **Sex:Temperature** | 0,00 | 0,00 | 1 | 57,63 | 0,41 | 5,27E-01 | 0,02 | 0,02 | 1 | 69,93 | 14,98 | 2,42E-04 |
| | **Odk** | **Sex** | 0,01 | 0,01 | 1 | 59,75 | 12,42 | 8,23E-04 | 0,04 | 0,04 | 1 | 69,58 | 34,76 | 1,21E-07 |
| | | **Temperature** | 0,10 | 0,10 | 1 | 59,45 | 92,03 | 1,10E-13 | 0,04 | 0,04 | 1 | 69,20 | 28,86 | 9,87E-07 |
| | | **Sex:Temperature** | 0,00 | 0,00 | 1 | 60,26 | 3,30 | 7,43E-02 | 0,01 | 0,01 | 1 | 69,94 | 10,51 | 1,82E-03 |
| | **Ran** | **Sex** | 0,04 | 0,04 | 1 | 58,01 | 18,82 | 5,81E-05 | 0,22 | 0,22 | 1 | 69,98 | 29,03 | 9,08E-07 |
| | | **Temperature** | 0,09 | 0,09 | 1 | 52,30 | 40,07 | 5,65E-08 | 0,06 | 0,06 | 1 | 69,75 | 8,38 | 5,06E-03 |
| | | **Sex:Temperature** | 0,00 | 0,00 | 1 | 26,61 | 0,05 | 8,30E-01 | 0,04 | 0,04 | 1 | 68,07 | 5,43 | 2,28E-02 |
| | **Cbk** | **Sex** | 0,01 | 0,01 | 1 | 61,00 | 0,63 | 4,31E-01 | 0,05 | 0,05 | 1 | 69,97 | 23,04 | 8,71E-06 |
| | | **Temperature** | 0,01 | 0,01 | 1 | 61,00 | 0,45 | 5,04E-01 | 0,00 | 0,00 | 1 | 69,72 | 0,68 | 4,11E-01 |
| | | **Sex:Temperature** | 0,03 | 0,03 | 1 | 61,00 | 2,06 | 1,56E-01 | 0,01 | 0,01 | 1 | 68,22 | 4,32 | 4,14E-02 |
| | **Cpa** | **Sex** | 1,09 | 1,09 | 1 | 59,93 | 6,89 | 1,10E-02 | 0,00 | 0,00 | 1 | 69,95 | 0,43 | 5,14E-01 |
| | | **Temperature** | 0,00 | 0,00 | 1 | 59,98 | 0,02 | 8,97E-01 | 0,01 | 0,01 | 1 | 69,68 | 2,48 | 1,19E-01 |
| | | **Sex:Temperature** | 0,06 | 0,06 | 1 | 45,53 | 0,38 | 5,39E-01 | 0,02 | 0,02 | 1 | 68,57 | 5,46 | 2,24E-02 |
| **OregonR** | **Pat** | **Sex** | 0,00 | 0,00 | 1 | 75,00 | 0,03 | 8,72E-01 | 0,49 | 0,49 | 1 | 32,94 | 212,38 | 6,42E-16 |
| | | **Temperature** | 0,05 | 0,05 | 1 | 75,00 | 2,10 | 1,52E-01 | 0,31 | 0,31 | 1 | 6,53 | 134,38 | 1,36E-05 |
| | | **Sex:Temperature** | 0,00 | 0,00 | 1 | 75,00 | 0,03 | 8,60E-01 | 0,00 | 0,00 | 1 | 8,05 | 0,11 | 7,51E-01 |
| | **Odk** | **Sex** | 0,00 | 0,00 | 1 | 37,60 | 0,00 | 9,62E-01 | 0,16 | 0,16 | 1 | 34,41 | 103,39 | 6,64E-12 |
| | | **Temperature** | 0,05 | 0,05 | 1 | 48,99 | 33,71 | 4,66E-07 | 0,00 | 0,00 | 1 | 12,27 | 0,15 | 7,02E-01 |
| | | **Sex:Temperature** | 0,04 | 0,04 | 1 | 20,33 | 25,91 | 5,33E-05 | 0,05 | 0,05 | 1 | 10,78 | 31,26 | 1,75E-04 |
| | **Ran** | **Sex** | 0,06 | 0,06 | 1 | 75,00 | 25,53 | 2,98E-06 | 0,22 | 0,22 | 1 | 37,58 | 23,12 | 2,47E-05 |
| | | **Temperature** | 0,13 | 0,13 | 1 | 75,00 | 53,46 | 2,41E-10 | 0,30 | 0,30 | 1 | 26,53 | 31,86 | 5,76E-06 |
| | | **Sex:Temperature** | 0,03 | 0,03 | 1 | 75,00 | 11,88 | 9,33E-04 | 0,54 | 0,54 | 1 | 15,57 | 56,50 | 1,46E-06 |
| | **Cbk** | **Sex** | 0,06 | 0,06 | 1 | 75,00 | 0,77 | 3,84E-01 | 0,02 | 0,02 | 1 | 42,20 | 20,72 | 4,47E-05 |
| | | **Temperature** | 0,00 | 0,00 | 1 | 75,00 | 0,03 | 8,73E-01 | 0,01 | 0,01 | 1 | 28,86 | 12,01 | 1,68E-03 |
| | | **Sex:Temperature** | 0,03 | 0,03 | 1 | 75,00 | 0,43 | 5,14E-01 | 0,00 | 0,00 | 1 | 18,09 | 0,07 | 7,98E-01 |
| | **Cpa** | **Sex** | 0,01 | 0,01 | 1 | 45,60 | 0,06 | 8,02E-01 | 0,05 | 0,05 | 1 | 44,08 | 17,31 | 1,45E-04 |
| | | **Temperature** | 0,00 | 0,00 | 1 | 66,43 | 0,00 | 9,85E-01 | 0,01 | 0,01 | 1 | 25,02 | 3,87 | 6,02E-02 |
| | | **Sex:Temperature** | 0,02 | 0,02 | 1 | 29,58 | 0,13 | 7,24E-01 | 0,00 | 0,00 | 1 | 17,97 | 0,31 | 5,85E-01 |

**Table S3. Windows of sensitivity for pigmentation plasticity in *D. melanogaster.*** Results of analysis of variance for the effect and interaction of strain and thermal regime on the different pigmentation traits per body part (model: *lm(Trait ~ Strain*Regime)*).

| | | Df | Sum Sq | Mean Sq | F value | Pr(>F) |
|---|---|---|---|---|---|---|
| Pat | Strain | 1 | 0,08 | 0,08 | 17,32 | 4,09E-05 |
| | Regime | 8 | 1,07 | 0,13 | 28,65 | <2,2e-16 |
| | Strain:Regime | 8 | 0,17 | 0,02 | 4,42 | 4,40E-05 |
| | Residuals | 312 | 1,46 | 0,00 | | |
| Odk | Strain | 1 | 0,00 | 0,00 | 1,02 | 3,13E-01 |
| | Regime | 8 | 1,25 | 0,16 | 40,31 | <2,2e-16 |
| | Strain:Regime | 8 | 0,22 | 0,03 | 6,96 | 1,63E-08 |
| | Residuals | 347 | 1,35 | 0,00 | | |
| Ran | Strain | 1 | 1,73 | 1,73 | 101,40 | <2,2e-16 |
| | Regime | 8 | 4,94 | 0,62 | 36,20 | <2,2e-16 |
| | Strain:Regime | 8 | 0,45 | 0,06 | 3,33 | 1,11E-03 |
| | Residuals | 347 | 5,92 | 0,02 | | |
| Cbk | Strain | 1 | 0,01 | 0,01 | 3,93 | 4,82E-02 |
| | Regime | 8 | 0,20 | 0,02 | 17,24 | <2,2e-16 |
| | Strain:Regime | 8 | 0,10 | 0,01 | 8,98 | 3,32E-11 |
| | Residuals | 347 | 0,49 | 0,00 | | |
| Cpa | Strain | 1 | 0,09 | 0,09 | 26,16 | 5,22E-07 |
| | Regime | 8 | 0,63 | 0,08 | 22,36 | <2,2e-16 |
| | Strain:Regime | 8 | 0,17 | 0,02 | 6,15 | 2,00E-07 |
| | Residuals | 347 | 1,23 | 0,00 | | |

**Table S4. Pigmentation components in *D. melanogaster* European populations.** Results of analysis of variance for the effect and interaction of location, genotype (i.e. isogenic line) and temperature on the different pigmentation traits per body part (model: *lm(Trait ~ Location * Location/Genotype * Temperature + (1|Replicate))*). Non-significant effects (p-value > 0.05) are shown in grey.

| | | THORAX | | | | | | ABDOMEN | | | | | |
|---|---|---|---|---|---|---|---|---|---|---|---|---|---|
| | | Sum Sq | Mean Sq | NumDF | DenDF | F value | Pr(>F) | Sum Sq | Mean Sq | NumDF | DenDF | F value | Pr(>F) |
| Pat | Location | 0,26 | 0,13 | 2 | 460,00 | 15,56 | 2,88E-07 | 0,00 | 0,00 | 2 | 467,79 | 0,51 | 6,02E-01 |
| | Temperature | 0,38 | 0,38 | 1 | 460,00 | 46,39 | 3,05E-11 | 2,34 | 2,34 | 1 | 428,39 | 655,94 | <2,2E-16 |
| | Location:Genotype | 1,13 | 0,09 | 12 | 460,00 | 11,42 | <2,2E-16 | 0,81 | 0,07 | 12 | 465,92 | 18,88 | <2,2E-16 |
| | Location:Temperature | 0,08 | 0,04 | 2 | 460,00 | 5,01 | 7,05E-03 | 0,03 | 0,02 | 2 | 467,79 | 4,34 | 1,36E-02 |
| | Location:Genotype:Temperature | 0,33 | 0,03 | 12 | 460,00 | 3,34 | 1,14E-04 | 0,16 | 0,01 | 12 | 465,78 | 3,82 | 1,39E-05 |
| Odk | Location | 0,39 | 0,19 | 2 | 458,59 | 83,57 | <2,2E-16 | 0,18 | 0,09 | 2 | 468,00 | 31,62 | 1,32E-13 |
| | Temperature | 0,13 | 0,13 | 1 | 457,75 | 58,55 | 1,18E-13 | 0,11 | 0,11 | 1 | 468,00 | 37,18 | 2,26E-09 |
| | Location:Genotype | 1,72 | 0,14 | 12 | 457,36 | 62,25 | <2,2E-16 | 0,96 | 0,08 | 12 | 468,00 | 28,20 | <2,2E-16 |
| | Location:Temperature | 0,15 | 0,07 | 2 | 458,59 | 31,66 | 1,32E-13 | 0,02 | 0,01 | 2 | 468,00 | 2,84 | 5,96E-02 |
| | Location:Genotype:Temperature | 0,33 | 0,03 | 12 | 457,35 | 11,82 | <2,2E-16 | 0,22 | 0,02 | 12 | 468,00 | 6,53 | 8,16E-11 |
| Ran | Location | 0,20 | 0,10 | 2 | 458,86 | 52,31 | <2,2E-16 | 0,57 | 0,29 | 2 | 464,52 | 35,87 | 3,25E-15 |
| | Temperature | 0,08 | 0,08 | 1 | 453,44 | 42,78 | 1,66E-10 | 0,38 | 0,38 | 1 | 359,23 | 47,39 | 2,61E-11 |
| | Location:Genotype | 0,27 | 0,02 | 12 | 457,32 | 12,14 | <2,2E-16 | 2,46 | 0,20 | 12 | 459,50 | 25,61 | <2,2E-16 |
| | Location:Temperature | 0,03 | 0,02 | 2 | 458,86 | 8,77 | 1,83E-04 | 0,06 | 0,03 | 2 | 464,52 | 4,02 | 1,85E-02 |
| | Location:Genotype:Temperature | 0,05 | 0,00 | 12 | 457,30 | 2,31 | 7,10E-03 | 0,62 | 0,05 | 12 | 459,60 | 6,45 | 1,21E-10 |
| Cbk | Location | 0,20 | 0,10 | 2 | 459,30 | 1,63 | 1,97E-01 | 0,11 | 0,05 | 2 | 468,18 | 10,05 | 5,32E-05 |
| | Temperature | 0,05 | 0,05 | 1 | 452,16 | 0,85 | 3,58E-01 | 0,11 | 0,11 | 1 | 460,04 | 20,82 | 6,47E-06 |
| | Location:Genotype | 2,00 | 0,17 | 12 | 458,86 | 2,70 | 1,54E-03 | 0,83 | 0,07 | 12 | 466,53 | 12,64 | <2,2E-16 |
| | Location:Temperature | 0,05 | 0,03 | 2 | 459,36 | 0,44 | 6,47E-01 | 0,04 | 0,02 | 2 | 468,18 | 3,95 | 2,00E-02 |
| | Location:Genotype:Temperature | 2,16 | 0,18 | 12 | 459,17 | 2,93 | 6,18E-04 | 0,26 | 0,02 | 12 | 466,45 | 4,03 | 5,61E-06 |
| Cpa | Location | 0,12 | 0,06 | 2 | 458,80 | 4,78 | 8,78E-03 | 0,05 | 0,02 | 2 | 467,51 | 6,20 | 2,20E-03 |
| | Temperature | 0,00 | 0,00 | 1 | 458,04 | 0,31 | 5,76E-01 | 0,24 | 0,24 | 1 | 467,92 | 60,79 | 4,16E-14 |
| | Location:Genotype | 1,29 | 0,11 | 12 | 457,74 | 8,56 | 9,71E-15 | 0,43 | 0,04 | 12 | 466,37 | 8,89 | 2,08E-15 |
| | Location:Temperature | 0,02 | 0,01 | 2 | 458,80 | 0,66 | 5,19E-01 | 0,03 | 0,01 | 2 | 467,51 | 3,18 | 4,26E-02 |
| | Location:Genotype:Temperature | 0,45 | 0,04 | 12 | 457,73 | 2,98 | 5,03E-04 | 0,18 | 0,01 | 12 | 466,33 | 3,74 | 1,99E-05 |

**Table S5. Pigmentation components in *Drosophila* species.** Results of analysis of variance for the effect and interaction of sex and temperature on the different pigmentation traits per body part (model: *lm(Trait ~ Sex * Temperature + (1|Replicate)*). Exceptionally for *D. simulans,* model: *lm(Trait ~ Strain * Sex * Temperature + (1|Replicate)*), where *Strain* corresponds to the different genetic backgrounds studied in this species (*D.sim* A and *D.sim* B). Non-significant effects (p-value > 0.05) are shown in grey.

| | | | THORAX | | | | | | ABDOMEN | | | | | |
|---|---|---|---|---|---|---|---|---|---|---|---|---|---|---|
| | | | Sum Sq | Mean Sq | NumDF | DenDF | F value | Pr(>F) | Sum Sq | Mean Sq | NumDF | DenDF | F value | Pr(>F) |
| *D.sim* | Pat | Strain | 0,02 | 0,02 | 1 | 106,00 | 1,46 | 2,30E-01 | 0,13 | 0,13 | 1 | 64,28 | 62,86 | 4,19E-11 |
| | | Sex | 0,29 | 0,29 | 1 | 106,00 | 19,97 | 1,98E-05 | 0,54 | 0,54 | 1 | 108,00 | 272,49 | <2,20E-16 |
| | | Temperature | 0,01 | 0,01 | 1 | 106,00 | 0,76 | 3,84E-01 | 0,50 | 0,50 | 1 | 106,54 | 252,49 | <2,20E-16 |
| | | Strain:Sex | 0,03 | 0,03 | 1 | 106,00 | 1,82 | 1,80E-01 | 0,00 | 0,00 | 1 | 108,00 | 0,55 | 4,59E-01 |
| | | Strain:Temperature | 0,07 | 0,07 | 1 | 106,00 | 4,77 | 3,11E-02 | 0,00 | 0,00 | 1 | 106,54 | 1,86 | 1,76E-01 |
| | | Sex:Temperature | 0,05 | 0,05 | 1 | 106,00 | 3,30 | 7,21E-02 | 0,03 | 0,03 | 1 | 107,19 | 15,94 | 1,20E-04 |
| | | Strain:Sex:Temperatur | 0,03 | 0,03 | 1 | 106,00 | 2,38 | 1,26E-01 | 0,00 | 0,00 | 1 | 107,19 | 0,45 | 5,03E-01 |
| | Odk | Strain | 0,01 | 0,01 | 1 | 105,92 | 3,79 | 5,43E-02 | 0,07 | 0,07 | 1 | 103,46 | 29,29 | 4,05E-07 |
| | | Sex | 0,08 | 0,08 | 1 | 104,47 | 54,68 | 3,68E-11 | 0,16 | 0,16 | 1 | 107,27 | 68,01 | 4,44E-13 |
| | | Temperature | 0,61 | 0,61 | 1 | 104,84 | 437,62 | <2,20E-16 | 0,65 | 0,65 | 1 | 107,89 | 267,48 | <2,20E-16 |
| | | Strain:Sex | 0,00 | 0,00 | 1 | 104,47 | 0,00 | 9,84E-01 | 0,01 | 0,01 | 1 | 107,27 | 3,07 | 8,28E-02 |
| | | Strain:Temperature | 0,00 | 0,00 | 1 | 104,84 | 0,56 | 4,55E-01 | 0,02 | 0,02 | 1 | 107,89 | 6,40 | 1,29E-02 |
| | | Sex:Temperature | 0,00 | 0,00 | 1 | 104,16 | 0,25 | 6,15E-01 | 0,04 | 0,04 | 1 | 106,58 | 16,21 | 1,06E-04 |
| | | Strain:Sex:Temperatur | 0,01 | 0,01 | 1 | 104,16 | 5,06 | 2,66E-02 | 0,01 | 0,01 | 1 | 106,58 | 2,37 | 1,26E-01 |
| | Ran | Strain | 0,00 | 0,00 | 1 | 33,51 | 4,52 | 4,10E-02 | 0,01 | 0,01 | 1 | 98,51 | 0,77 | 3,82E-01 |
| | | Sex | 0,00 | 0,00 | 1 | 105,17 | 1,33 | 2,51E-01 | 0,37 | 0,37 | 1 | 107,52 | 38,17 | 1,18E-08 |
| | | Temperature | 0,03 | 0,03 | 1 | 101,69 | 40,40 | 5,95E-09 | 0,23 | 0,23 | 1 | 108,00 | 22,95 | 5,33E-06 |
| | | Strain:Sex | 0,00 | 0,00 | 1 | 105,17 | 5,69 | 1,89E-02 | 0,07 | 0,07 | 1 | 107,52 | 6,71 | 1,09E-02 |
| | | Strain:Temperature | 0,01 | 0,01 | 1 | 101,69 | 10,98 | 1,28E-03 | 0,27 | 0,27 | 1 | 108,00 | 27,22 | 8,83E-07 |
| | | Sex:Temperature | 0,00 | 0,00 | 1 | 105,47 | 0,47 | 4,96E-01 | 0,17 | 0,17 | 1 | 106,74 | 17,79 | 5,19E-05 |
| | | Strain:Sex:Temperatur | 0,01 | 0,01 | 1 | 105,47 | 17,51 | 5,93E-05 | 0,13 | 0,13 | 1 | 106,74 | 13,13 | 4,48E-04 |
| | Cbk | Strain | 0,29 | 0,29 | 1 | 106,00 | 5,41 | 2,20E-02 | 0,01 | 0,01 | 1 | 107,92 | 7,14 | 8,70E-03 |
| | | Sex | 0,09 | 0,09 | 1 | 106,00 | 1,74 | 1,90E-01 | 0,17 | 0,17 | 1 | 106,47 | 125,72 | <2,20E-16 |
| | | Temperature | 0,02 | 0,02 | 1 | 106,00 | 0,29 | 5,93E-01 | 0,00 | 0,00 | 1 | 106,89 | 1,12 | 2,92E-01 |
| | | Strain:Sex | 0,58 | 0,58 | 1 | 106,00 | 10,93 | 1,29E-03 | 0,00 | 0,00 | 1 | 106,47 | 2,77 | 9,89E-02 |
| | | Strain:Temperature | 0,28 | 0,28 | 1 | 106,00 | 5,23 | 2,42E-02 | 0,05 | 0,05 | 1 | 106,89 | 34,93 | 4,13E-08 |
| | | Sex:Temperature | 0,13 | 0,13 | 1 | 106,00 | 2,41 | 1,23E-01 | 0,03 | 0,03 | 1 | 106,19 | 25,69 | 1,71E-06 |
| | | Strain:Sex:Temperatur | 0,04 | 0,04 | 1 | 106,00 | 0,77 | 3,81E-01 | 0,00 | 0,00 | 1 | 106,19 | 3,54 | 6,27E-02 |
| | Cpa | Strain | 0,33 | 0,33 | 1 | 77,57 | 3,50 | 6,51E-02 | 0,10 | 0,10 | 1 | 41,27 | 34,47 | 6,45E-07 |
| | | Sex | 0,01 | 0,01 | 1 | 105,92 | 0,13 | 7,16E-01 | 0,16 | 0,16 | 1 | 107,59 | 55,86 | 2,19E-11 |
| | | Temperature | 1,16 | 1,16 | 1 | 105,72 | 12,21 | 6,94E-04 | 0,07 | 0,07 | 1 | 103,58 | 24,11 | 3,41E-06 |
| | | Strain:Sex | 0,00 | 0,00 | 1 | 105,92 | 0,00 | 9,65E-01 | 0,03 | 0,03 | 1 | 107,59 | 8,96 | 3,42E-03 |
| | | Strain:Temperature | 0,14 | 0,14 | 1 | 105,72 | 1,42 | 2,35E-01 | 0,00 | 0,00 | 1 | 103,58 | 0,06 | 8,13E-01 |
| | | Sex:Temperature | 0,30 | 0,30 | 1 | 104,83 | 3,18 | 7,74E-02 | 0,00 | 0,00 | 1 | 107,52 | 0,86 | 3,55E-01 |
| | | Strain:Sex:Temperatur | 0,26 | 0,26 | 1 | 104,83 | 2,76 | 9,99E-02 | 0,01 | 0,01 | 1 | 107,52 | 2,03 | 1,57E-01 |

Continuation **Table S5**

| | | | THORAX | | | | | | ABDOMEN | | | | | |
|---|---|---|---|---|---|---|---|---|---|---|---|---|---|---|
| | | | Sum Sq | Mean Sq | NumDF | DenDF | F value | Pr(>F) | Sum Sq | Mean Sq | NumDF | DenDF | F value | Pr(>F) |
| *D.mal* | **Pat** | **Sex** | 0,08 | 0,08 | 1 | 56,99 | 6,95 | 1,08E-02 | 0,13 | 0,13 | 1 | 57,00 | 24,09 | 8,08E-06 |
| | | **Temperature** | 0,23 | 0,23 | 1 | 54,08 | 20,61 | 3,18E-05 | 0,25 | 0,25 | 1 | 57,00 | 45,48 | 8,49E-09 |
| | | **Sex:Temperature** | 0,00 | 0,00 | 1 | 56,60 | 0,23 | 6,35E-01 | 0,00 | 0,00 | 1 | 57,00 | 0,90 | 3,46E-01 |
| | **Odk** | **Sex** | 0,00 | 0,00 | 1 | 53,39 | 0,12 | 7,34E-01 | 0,17 | 0,17 | 1 | 56,86 | 34,80 | 2,12E-07 |
| | | **Temperature** | 0,02 | 0,02 | 1 | 53,88 | 5,58 | 2,19E-02 | 0,00 | 0,00 | 1 | 55,94 | 0,00 | 9,90E-01 |
| | | **Sex:Temperature** | 0,01 | 0,01 | 1 | 53,34 | 4,60 | 3,65E-02 | 0,01 | 0,01 | 1 | 56,52 | 1,11 | 2,96E-01 |
| | **Ran** | **Sex** | 0,00 | 0,00 | 1 | 57,00 | 2,12 | 1,50E-01 | 0,04 | 0,04 | 1 | 55,94 | 4,28 | 4,31E-02 |
| | | **Temperature** | 0,02 | 0,02 | 1 | 55,04 | 10,77 | 1,80E-03 | 0,31 | 0,31 | 1 | 56,62 | 30,35 | 9,18E-07 |
| | | **Sex:Temperature** | 0,00 | 0,00 | 1 | 56,84 | 0,22 | 6,38E-01 | 0,28 | 0,28 | 1 | 55,33 | 27,49 | 2,56E-06 |
| | **Cbk** | **Sex** | 0,03 | 0,03 | 1 | 57,00 | 0,64 | 4,26E-01 | 0,03 | 0,03 | 1 | 55,75 | 26,06 | 4,14E-06 |
| | | **Temperature** | 0,14 | 0,14 | 1 | 57,00 | 2,92 | 9,28E-02 | 0,00 | 0,00 | 1 | 56,79 | 1,27 | 2,65E-01 |
| | | **Sex:Temperature** | 0,05 | 0,05 | 1 | 57,00 | 1,02 | 3,16E-01 | 0,00 | 0,00 | 1 | 55,58 | 1,26 | 2,66E-01 |
| | **Cpa** | **Sex** | 0,00 | 0,00 | 1 | 57,00 | 0,01 | 9,21E-01 | 0,08 | 0,08 | 1 | 56,26 | 13,16 | 6,18E-04 |
| | | **Temperature** | 0,02 | 0,02 | 1 | 57,00 | 0,35 | 5,56E-01 | 0,00 | 0,00 | 1 | 56,91 | 0,00 | 9,69E-01 |
| | | **Sex:Temperature** | 0,00 | 0,00 | 1 | 57,00 | 0,08 | 7,74E-01 | 0,00 | 0,00 | 1 | 55,93 | 0,40 | 5,28E-01 |
| *D.mojbaj* | **Pat** | **Sex** | 0,03 | 0,03 | 1 | 82,00 | 4,57 | 3,56E-02 | 0,07 | 0,07 | 1 | 79,95 | 31,44 | 2,84E-07 |
| | | **Temperature** | 0,97 | 0,97 | 1 | 82,00 | 129,47 | <2,00E-16 | 0,39 | 0,39 | 1 | 66,09 | 183,19 | <2,20E-16 |
| | | **Sex:Temperature** | 0,02 | 0,02 | 1 | 82,00 | 2,20 | 1,42E-01 | 0,00 | 0,00 | 1 | 79,99 | 0,17 | 6,80E-01 |
| | **Odk** | **Sex** | 0,01 | 0,01 | 1 | 82,00 | 3,93 | 5,07E-02 | 0,04 | 0,04 | 1 | 79,96 | 7,30 | 8,41E-03 |
| | | **Temperature** | 0,47 | 0,47 | 1 | 82,00 | 226,13 | <2,00E-16 | 0,25 | 0,25 | 1 | 71,04 | 47,44 | 1,88E-09 |
| | | **Sex:Temperature** | 0,01 | 0,01 | 1 | 82,00 | 4,22 | 4,31E-02 | 0,05 | 0,05 | 1 | 79,81 | 9,13 | 3,38E-03 |
| | **Ran** | **Sex** | 0,01 | 0,01 | 1 | 81,77 | 6,31 | 1,40E-02 | 0,02 | 0,02 | 1 | 79,93 | 1,37 | 2,45E-01 |
| | | **Temperature** | 0,13 | 0,13 | 1 | 73,50 | 81,17 | 1,69E-13 | 0,22 | 0,22 | 1 | 50,76 | 14,05 | 4,56E-04 |
| | | **Sex:Temperature** | 0,02 | 0,02 | 1 | 81,77 | 15,23 | 1,94E-04 | 0,27 | 0,27 | 1 | 80,00 | 17,43 | 7,54E-05 |
| | **Cbk** | **Sex** | 0,00 | 0,00 | 1 | 82,00 | 0,43 | 5,12E-01 | 0,05 | 0,05 | 1 | 79,99 | 11,03 | 1,35E-03 |
| | | **Temperature** | 0,70 | 0,70 | 1 | 82,00 | 90,14 | 7,49E-15 | 0,04 | 0,04 | 1 | 65,86 | 10,21 | 2,15E-03 |
| | | **Sex:Temperature** | 0,01 | 0,01 | 1 | 82,00 | 1,22 | 2,73E-01 | 0,00 | 0,00 | 1 | 80,00 | 0,02 | 8,95E-01 |
| | **Cpa** | **Sex** | 0,01 | 0,01 | 1 | 81,72 | 1,92 | 1,69E-01 | 0,18 | 0,18 | 1 | 79,14 | 32,02 | 2,35E-07 |
| | | **Temperature** | 0,67 | 0,67 | 1 | 74,17 | 107,59 | 4,36E-16 | 0,01 | 0,01 | 1 | 79,14 | 1,29 | 2,60E-01 |
| | | **Sex:Temperature** | 0,00 | 0,00 | 1 | 81,70 | 0,00 | 9,59E-01 | 0,07 | 0,07 | 1 | 78,96 | 11,58 | 1,05E-03 |
| *D.mojmoj* | **Pat** | **Sex** | 0,00 | 0,00 | 1 | 23,98 | 0,12 | 7,35E-01 | 0,04 | 0,04 | 1 | 23,02 | 21,50 | 1,15E-04 |
| | | **Temperature** | 0,11 | 0,11 | 1 | 18,17 | 10,03 | 5,29E-03 | 0,36 | 0,36 | 1 | 23,87 | 194,32 | 5,81E-13 |
| | | **Sex:Temperature** | 0,00 | 0,00 | 1 | 23,98 | 0,04 | 8,40E-01 | 0,01 | 0,01 | 1 | 23,02 | 2,89 | 1,03E-01 |
| | **Odk** | **Sex** | 0,00 | 0,00 | 1 | 23,55 | 1,47 | 2,38E-01 | 0,04 | 0,04 | 1 | 23,91 | 14,21 | 9,45E-04 |
| | | **Temperature** | 0,32 | 0,32 | 1 | 5,88 | 452,42 | 8,74E-07 | 0,12 | 0,12 | 1 | 20,88 | 47,59 | 8,38E-07 |
| | | **Sex:Temperature** | 0,00 | 0,00 | 1 | 23,55 | 1,72 | 2,03E-01 | 0,00 | 0,00 | 1 | 23,91 | 0,06 | 8,16E-01 |
| | **Ran** | **Sex** | 0,00 | 0,00 | 1 | 24,00 | 0,00 | 9,91E-01 | 0,00 | 0,00 | 1 | 23,86 | 0,12 | 7,29E-01 |
| | | **Temperature** | 0,07 | 0,07 | 1 | 24,00 | 180,37 | 1,17E-12 | 0,00 | 0,00 | 1 | 10,33 | 0,02 | 9,02E-01 |
| | | **Sex:Temperature** | 0,00 | 0,00 | 1 | 24,00 | 0,13 | 7,22E-01 | 0,13 | 0,13 | 1 | 23,86 | 12,28 | 1,84E-03 |
| | **Cbk** | **Sex** | 0,01 | 0,01 | 1 | 23,00 | 0,85 | 3,66E-01 | 0,00 | 0,00 | 1 | 24,00 | 0,19 | 6,69E-01 |
| | | **Temperature** | 0,12 | 0,12 | 1 | 3,01 | 11,53 | 4,24E-02 | 0,00 | 0,00 | 1 | 24,00 | 0,18 | 6,72E-01 |
| | | **Sex:Temperature** | 0,02 | 0,02 | 1 | 23,00 | 1,50 | 2,33E-01 | 0,02 | 0,02 | 1 | 24,00 | 7,35 | 1,22E-02 |
| | **Cpa** | **Sex** | 0,04 | 0,04 | 1 | 22,86 | 3,47 | 7,54E-02 | 0,00 | 0,00 | 1 | 24,00 | 0,90 | 3,53E-01 |
| | | **Temperature** | 0,19 | 0,19 | 1 | 23,84 | 16,78 | 4,18E-04 | 0,01 | 0,01 | 1 | 24,00 | 2,14 | 1,56E-01 |
| | | **Sex:Temperature** | 0,09 | 0,09 | 1 | 22,86 | 8,15 | 9,00E-03 | 0,00 | 0,00 | 1 | 24,00 | 0,11 | 7,44E-01 |

Continuation **Table S5**

| | | | THORAX | | | | | | ABDOMEN | | | | | |
|---|---|---|---|---|---|---|---|---|---|---|---|---|---|---|
| | | | Sum Sq | Mean Sq | NumDF | DenDF | F value | Pr(>F) | Sum Sq | Mean Sq | NumDF | DenDF | F value | Pr(>F) |
| *D.rep* | Pat | Sex | 0,00 | 0,00 | 1 | 49,00 | 0,36 | 5,54E-01 | 0,00 | 0,00 | 1 | 46,87 | 2,02 | 1,62E-01 |
| | | Temperature | 0,04 | 0,04 | 1 | 49,00 | 5,95 | 1,84E-02 | 0,00 | 0,00 | 1 | 48,34 | 2,19 | 1,45E-01 |
| | | Sex:Temperature | 0,00 | 0,00 | 1 | 49,00 | 0,00 | 9,84E-01 | 0,00 | 0,00 | 1 | 46,02 | 0,47 | 4,97E-01 |
| | Odk | Sex | 0,00 | 0,00 | 1 | 45,37 | 0,09 | 7,70E-01 | 0,07 | 0,07 | 1 | 49,00 | 19,57 | 5,39E-05 |
| | | Temperature | 0,07 | 0,07 | 1 | 48,26 | 50,25 | 5,24E-09 | 0,00 | 0,00 | 1 | 49,00 | 0,11 | 7,40E-01 |
| | | Sex:Temperature | 0,00 | 0,00 | 1 | 44,19 | 1,30 | 2,60E-01 | 0,01 | 0,01 | 1 | 49,00 | 3,90 | 5,40E-02 |
| | Ran | Sex | 0,00 | 0,00 | 1 | 48,96 | 2,31 | 1,35E-01 | 0,14 | 0,14 | 1 | 49,00 | 8,97 | 4,29E-03 |
| | | Temperature | 0,04 | 0,04 | 1 | 46,59 | 87,75 | 2,75E-12 | 0,00 | 0,00 | 1 | 49,00 | 0,00 | 9,55E-01 |
| | | Sex:Temperature | 0,00 | 0,00 | 1 | 47,62 | 2,48 | 1,22E-01 | 0,13 | 0,13 | 1 | 49,00 | 8,48 | 5,38E-03 |
| | Cbk | Sex | 0,06 | 0,06 | 1 | 44,89 | 7,75 | 7,85E-03 | 0,01 | 0,01 | 1 | 48,70 | 6,80 | 1,21E-02 |
| | | Temperature | 0,22 | 0,22 | 1 | 48,09 | 29,38 | 1,89E-06 | 0,00 | 0,00 | 1 | 46,11 | 0,88 | 3,53E-01 |
| | | Sex:Temperature | 0,16 | 0,16 | 1 | 44,61 | 20,68 | 4,13E-05 | 0,00 | 0,00 | 1 | 46,99 | 0,44 | 5,10E-01 |
| | Cpa | Sex | 0,03 | 0,03 | 1 | 49,00 | 0,98 | 3,26E-01 | 0,00 | 0,00 | 1 | 47,02 | 0,01 | 9,27E-01 |
| | | Temperature | 0,68 | 0,68 | 1 | 45,04 | 22,07 | 2,50E-05 | 0,00 | 0,00 | 1 | 46,46 | 0,10 | 7,56E-01 |
| | | Sex:Temperature | 0,00 | 0,00 | 1 | 46,86 | 0,03 | 8,53E-01 | 0,01 | 0,01 | 1 | 45,29 | 1,83 | 1,83E-01 |
